# Subtle Pupil-Size Changes Associated With Exploration Do Not Affect Visual Sensitivity

**DOI:** 10.64898/2026.06.05.730459

**Authors:** Ward Claeys, Veera Ruuskanen, Sebastiaan Mathôt

**Affiliations:** Department of Experimental Psychology, Ghent University, Belgium; Department of Psychology, University of Groningen, The Netherlands

## Abstract

When we feel restless and easily distracted, continuously switching tasks (*exploration*), our pupils tend to be large. In contrast, when we are calmly focused on a single task (*exploitation*), our pupils tend to be small. According to the Adaptive Gain Theory (AGT), a switch from exploitation to exploration is associated with an increase in norepinephrine in the locus coeruleus, which in turn triggers pupil dilation. However, the AGT does not provide a functional explanation of why exploration triggers pupil dilation. One possibility is that visual sensitivity, which increases with pupil size, is especially important during exploration. We set out to provide evidence consistent with this functional explanation, as well as to replicate two key previous results. Participants performed a four-armed bandit task, which induces both exploration and exploitation behavior. During the task, participants also needed to detect an occasional and unpredictable near-threshold peripheral flash. We replicated two key results: pupils were larger during exploration than during exploitation; and increased pupil size (overall, independent of exploration status) was associated with increased visual sensitivity. However, most importantly, we did *not* find that visual sensitivity was higher during exploration than during exploitation; probably, the reliable-yet-tiny increase in pupil size during exploration was too small to affect visual sensitivity. We conclude that key previous results are replicable; however, common experimental paradigms, such as the four-armed bandit task, induce only small changes in exploration behavior. Therefore, more powerful paradigms are required in order to test functional explanations of pupil-size changes during exploration and exploitation.

---

Should I go to dinner at my favorite restaurant or should I try a different one? This is an example of an exploration-exploitation dilemma. When you know a restaurant is good, you can continue to go to that same restaurant, which would be an instance of exploitation. (In daily life, the term exploitation often has the negative connotation of taking advantage of someone. However, in this context, it is a neutral term that refers to continuing to do what you were doing.) However, this means you will never find an even better restaurant, because this would require trying out new restaurants, which would be an instance of exploration. This dilemma is not merely present when looking for restaurants, but is everywhere: deciding whether to keep doing research on the same topic or to move to a different topic; deciding whether to get the same coffee each day or to try something different; all of these, and many more, are examples of deciding between exploitation and exploration.

The difference between exploration and exploitation modes of behavior is central to the Adaptive Gain Theory (AGT; Aston-Jones & Cohen, 2005). Exploitation is characterized by focus on a single task, while exploration is characterized by distractibility and frequent task-switching. The AGT makes specific assumptions about the underlying processes, and puts the locus coeruleus norepinephrine (LC-NE) system forward as the main driver of the exploration-exploitation trade-off. The LC is a nucleus located in the brainstem which supplies the majority of the NE available to the brain, thus mediating arousal. This theory is based on studies that found positive correlations between the tendency to engage in exploration behavior on the one hand, and norepinephrine levels and LC activity on the other hand (Cohen et al., 2007; Doya, 2002; Gilzenrat et al., 2010; Wilson et al., 2021). LC activity and NE levels are also themselves positively correlated (Aston-Jones & Cohen, 2005; Benarroch, 2009; Berridge & Waterhouse, 2003; Grzanna & Molliver, 1980).

LC activity and NE levels are difficult to measure, because doing so requires invasive procedures, due to the location of the LC. However, research has shown that these both correlate with pupil size (Costa & Rudebeck, 2016; Gilzenrat et al., 2010; Joshi et al., 2016; Murphy et al., 2014; Reimer et al., 2016). That is, large pupils are associated with high levels of tonic LC activity and NE, whereas small pupils, with occasional brief dilations, are associated with low levels of tonic LC activity and NE, with occasional phasic bursts. (Here, tonic activity refers to sustained activity, whereas phasic activity refers to brief bursts of activity.) Based on this, many researchers have used pupil size as a non-invasive proxy for the LC-NE system in order to test predictions of the AGT (Gilzenrat et al., 2010; Hayes & Petrov, 2016; Jepma & Nieuwenhuis, 2011; but see Ruuskanen & Mathôt, 2026a for a critical review).

For example, in a landmark study by Jepma and Nieuwenhuis (2011), participants were asked to complete a four-armed bandit task (explained in depth by Daw et al., 2006) to investigate how pupil size differs depending on whether participants engage in exploration or exploitation. This task is a gambling game in which four slot machines provide points, ranging from 1 to 100 (points changed gradually and independently for all machines). Expected payoffs and choices were modeled for each participant. Subsequently, the choices of the participants were compared to the choices of the model. If the choice of the participant was in line with the choice of the model, the trial was labeled as exploitation; if the choice of the participant was different from the choice of the model, the trial was labeled as exploration. The authors found that the participants’ pupils were larger before they made exploration choices as opposed to exploitation choices.

In line with the LC-NE system’s role in arousal regulation, research has shown that pupil size correlates positively with arousal. In a seminal study by Hess and Polt (1960), participants were asked to view interesting, emotional, or neutral pictures while their pupil size was tracked. They found that pupils dilated more in response to interesting and emotional pictures than to neutral pictures. Later studies found that pupil dilation was present for both pleasant and unpleasant arousing stimuli (Bradley et al., 2008). That is, arousal (intensity) and not valence (positive vs negative), was the main driver of pupil dilation. More generally, research into the effect of arousal and cognitive factors on pupil dilation has concluded that anything that activates the mind has an influence on pupil dilation (Beatty & Lucero-Wagoner, 2000; Loewenfeld, 1958; Mathôt, 2018).

Furthermore, a distinct-but-related line of research has focused on the relationship between pupil size and attentional breadth. The general result from these studies is that pupil size increases when participants covertly attend to the periphery as opposed to the central part of the visual field (Brocher et al., 2018; Daniels et al., 2012; Klatt et al., 2021; Kolnes et al., 2024; Vilotijević & Mathôt, 2023).

The aforementioned pupil-size changes associated with exploration, arousal, and attentional breadth are often treated as distinct processes. However, they may all reflect similar underlying mechanisms. A possible, functional explanation relates to how pupil size affects visual perception (reviewed in Vilotijević & Mathôt, 2024). Specifically, when the pupil dilates, more light enters the eye, which results in increased visual sensitivity: the ability to detect faint stimuli. Conversely, when the pupil constricts, light that enters the eye is focused better, which results in increased visual acuity: the ability to discriminate fine detail.

Many studies have confirmed the hypothesis that changes in pupil size result in behaviorally relevant changes in visual sensitivity and acuity (Campbell & Gregory, 1960; Eberhardt et al., 2022; Mathôt & Ivanov, 2019; Wang et al., 2021; Woodhouse, 1975). In a study by Mathôt and Ivanov (2019), pupil size was manipulated by changing the brightness of the visual periphery, resulting in large pupils (dark periphery) or small pupils (bright periphery). Task-relevant stimuli were presented on a central gray disk that remained constant in luminance throughout the entire experiment. Both a discrimination and a detection task were performed by the participants. The discrimination task required discriminating the orientation of a Gabor patch (a patch of oriented lines) that was presented in the center of the screen, while the detection task required detecting the presence of a Gabor patch that sometimes appeared at a random location. Results indicated a positive relationship between pupil size and visual sensitivity on the detection task. However, there was no relationship between pupil size and visual acuity on the discrimination task. This latter null result was likely due to ceiling effects, because in a follow-up experiment with smaller stimuli, the authors did find a clear negative relationship between pupil size and visual acuity. The association between visual sensitivity and pupil dilation was later conceptually replicated in a study by Ruuskanen, Boehler, and Mathôt (2025; see also Ruuskanen & Mathôt, 2026b), who investigated the effect of pupil size on detection of near-threshold stimuli. They again found that larger pupils allowed participants to better detect visual near-threshold stimuli. This effect was found within the range of pupil-size fluctuations that naturally occur. Taken together, these results show that visual sensitivity is highest for large pupils, while visual acuity is highest for small pupils.

As the studies reviewed above have shown, pupil-size changes affect both detection and acuity, but in different ways. Pupil constriction increases visual acuity and is mainly relevant in tasks where participants have to focus tasks in central vision. In contrast, pupil dilation increases sensitivity and is mainly relevant in tasks where participants have to be vigilant in order to detect targets in unpredictable locations. As discussed, what exploration, increased arousal and increased attentional breadth potentially have in common is this emphasis on vigilance and peripheral vision (Mathôt, 2020; Vilotijević & Mathôt, 2024). And through this emphasis, they may all be associated with pupil dilation.

Taken together, the goal of the current study is twofold. The first goal is to replicate two key results: the finding of Jepma and Nieuwenhuis (2011) that pupils are larger during exploration than during exploitation; and the finding by Ruuskanen, Boehler, and Mathôt (2025; see also Ruuskanen & Mathôt, 2026b) that visual sensitivity is higher for large pupils than for small pupils. The second goal is to connect these two studies by testing whether pupil dilation during exploration is associated with increased visual sensitivity. If so, this would offer a functional explanation for the link between pupil dilation and exploration behavior as posited by the AGT.

## Materials and methods

### Materials and availability

All experimental materials are available from https://osf.io/8zd4g/.

### Participants

The sample consisted of 31 participants. Participants were undergraduate psychology students who received partial course credits for participating. Additionally, participants received a chocolate bar after the study. All participants had normal vision and did not wear glasses or contact lenses. Sample size was chosen to be considerably higher than in previous studies (Jepma & Nieuwenhuis, 2011, N = 17; Ruuskanen & Mathôt, 2026b, N = 19) and followed our lab’s rule of thumb to use a sample size of 30 participants when no a priori effect size can be estimated (one additional participant was tested due to over-registration).

Based on a list of criteria developed by the Ethics committee at the psychology department of the University of Groningen, the study was exempt from full ethical review (study code: PSY-2425-S-0075). All participants provided written informed consent prior to participation.

### Task

The paradigm is based on the design by Jepma and Nieuwenhuis (2011) combined with the paradigm of Ruuskanen et al. (2025). It consisted of two phases. The first phase served to calibrate, through a staircase procedure, the difficulty of a visual-detection task. In this first phase, four slot machines, as in a four-armed bandit task (see Fig. 1a), were presented on screen, although these were not (yet) task relevant. Slot machines had different colors but were roughly isoluminant. These slots stayed on screen whilst, occasionally, a white flash, which was a white luminance patch with a Gaussian envelope, was presented for 50 ms in the periphery of participants’ visual field at a random location with a distance 10.97° from the central fixation point. Participants pressed the spacebar whenever they detected a flash. Through a 3-up-1-down staircase procedure, target contrast decreased or increased in order to fix the accuracy of the participants to 75%. This first phase consisted of 120 trials.

**Figure 1.**
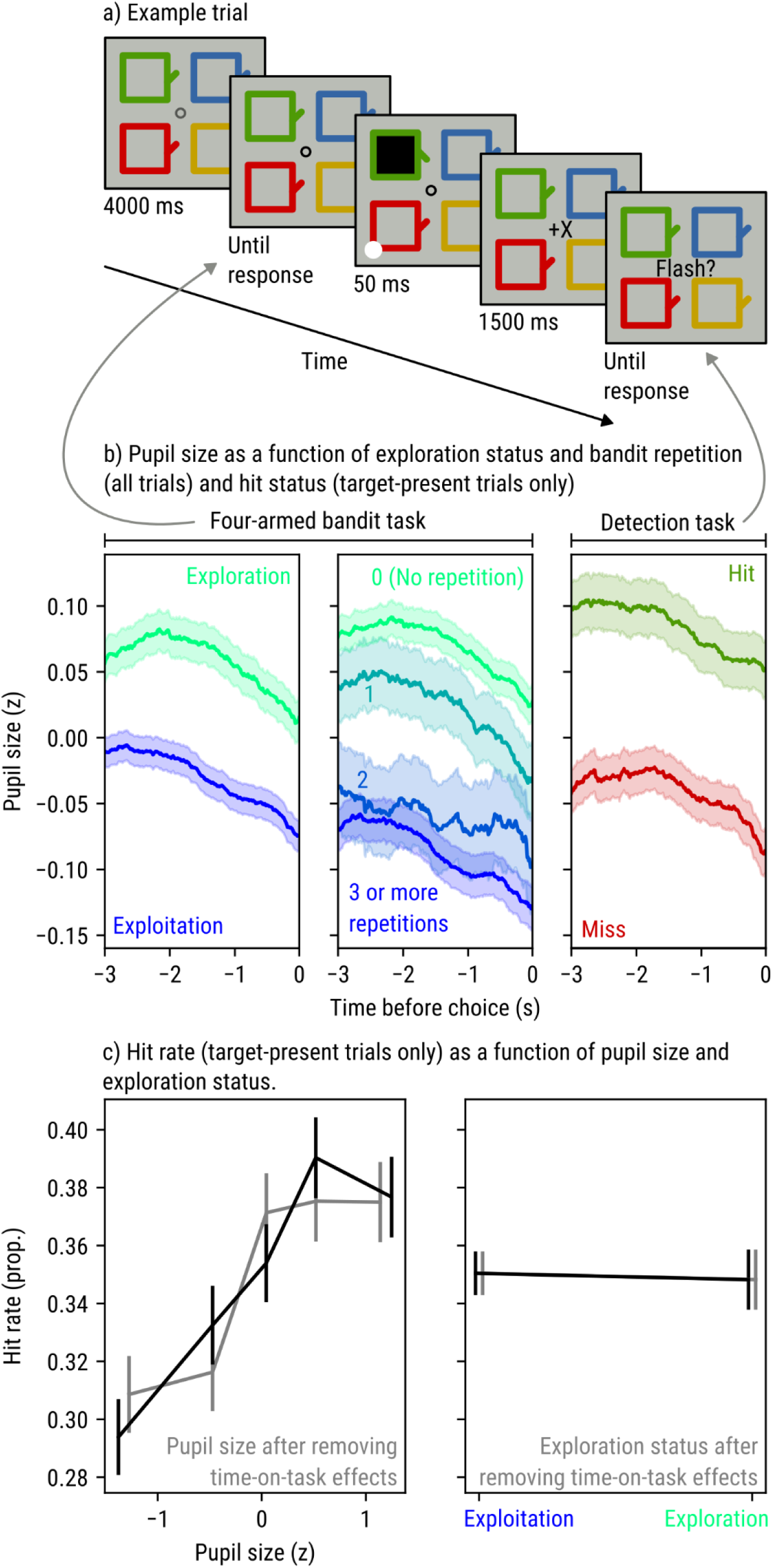
Overview of experimental procedure and main results. a) A schematic example of a target-present trial in which a white stimulus is briefly flashed after the selection of the green bandit. b) Pupil size as a function of exploration status and bandit repetition, which reflect the response on the four-armed bandit task, and hit status, which reflects behavior on target-present trials of the detection task. c) Hit rate (target-present trials only) as a function of pupil size and exploration status. Gray lines correspond to residual pupil-size and exploration values after removing time-on-task effects.

After this staircase procedure, the actual bandit task started. Again, the four slot machines appeared on screen, but this time they were task relevant. After a delay of 4 s, the initially gray fixation dot turned black, indicating to participants that they should choose a slot machine by using the keys “f”, “v”, “j” and “n”. The chosen slot would spin upon selection and a certain number of points were given to the participants. The four different slot machines differed in the (average) number of points that participants received when choosing them. This average changed gradually and independently from trial to trial, such that slot machines gradually became more or less profitable over time, without this being directly visible to the participants. The machines provided points between 1 and 100, drawn from a Gaussian distribution with a gradually changing mean and standard deviation of 4. The current mean was updated using a decaying Gaussian random walk procedure with decay parameter of 0.9836, decay center of 50 and diffusion noise of 2.8 (as in Daw et al., 2006).

On half of the trials, at the moment that participants selected a machine, a white flash was presented for 50ms in the periphery of the participants’ visual field, in the exact same way as in the first phase of the experiment. On every trial, after receiving their points, a message appeared asking the participant if they had seen a flash. Participants responded with the space-bar upon detection of a flash (Fig. 1a).

In total, the second phase contained 470 trials (trial duration depended on response speed, but maximum trial length was 8.05 seconds), split into 9 blocks of 50 trials and one practice block of 20 trials. Practice trials were included to ensure participants understood the task. Both during the practice trials and in the 50 trials of block 5, the staircase procedure on the responses to the flashes (as in the first phase of the experiment) ran again to ensure a 75% accuracy. Both the practice block and the staircase block in the middle of the experiment (block 5) were excluded from analysis. In other words, only blocks during which the difficulty level was fixed (i.e., the staircase was not running) were included in the analysis.

During the second phase of the experiment, pupil size was measured. To avoid unwanted pupil-size changes due to visual changes or eye movements (Knapen et al., 2016; Mathôt et al., 2015), the four bandits and a central fixation cross were presented throughout the entire second phase. For the same reason, and different from the study by Jepma and Nieuwenhuis (2011), points were presented in the middle of the screen.

In the instructions, participants were told that if they beat the high score of the experimenter on the bandit task, they would receive a chocolate bar (to encourage participants). However, as all values were randomized, the possibility of beating this high score was not ensured. Therefore, all participants received a chocolate bar after the experiment.

The experiment and its stimuli were created with and controlled by OpenSesame (version 3.3.9, Lentiform Loewenfeld) (Mathôt et al., 2012).

### Pupil size measurement

Pupil size was measured using an Eyelink 1000 eye tracker. Before starting the task, a 5-point calibration procedure was performed and a drift correction was performed at the beginning of each block (except block 5). Pupil size was measured during all trials in the second phase of the experiment, but not the first (staircase phase) because in this phase pupil size was not of interest. Participants rested their head on a chinrest throughout the entire experiment, keeping the distance to the eyetracker fixed at roughly 60 cm.

### Procedure

Participants arrived in the lab and signed an informed consent after receiving information about the study. Task instructions were provided verbally before starting the task and on the screen at the beginning of the task. The eye tracker was calibrated and participants performed the task. Participants were tested in a dimly lit room (6 lux). The whole session took approximately 1 hour per participant.

### Preprocessing

Pupil size data was parsed using EyelinkParser (Mathôt & Vilotijević, 2022). Data was downsampled to 100 Hz. Blinks were interpolated with cubic spline interpolation if possible, and with linear interpolation otherwise. Samples were rejected whenever the pupil was lost for longer than 500 ms, as this is unlikely to be a blink. Finally, pupil traces were z-scored separately per participant.

Trials were excluded when participants did not respond to the bandit selection, or when no pupil signal was recorded. No participants were excluded.

### Results Classification of exploration vs exploitation behavior

On each trial, the expected payoff was determined for each bandit using a mean-tracking algorithm (a Kalman filter) with the same parameters as used by Jepma and Nieuwenhuis (2011). Trials on which the bandit with the highest expected payoff was selected were labeled exploitation trials; other trials were labeled exploration trials. The proportion of exploration trials per participant ranged from 18% to 63% with a mean of 35%.

### Pupil-size window of interest

We analyzed pupil size during the three seconds before participants were prompted to select a bandit. This ensured that pupil size did not depend on the response time of the participant. All analyses are performed on the trial-level averages of pupil size during this time window.

### Time-on-task effects

All variables of interest vary systematically over the course of the experiment as well as over individual blocks (Fig. 2). Specifically, pupil size, exploration rate, and hit rate all decrease over the course of a block. Pupil size and exploration rate also decrease slightly over the course of the full experiment.

**Figure 2.**
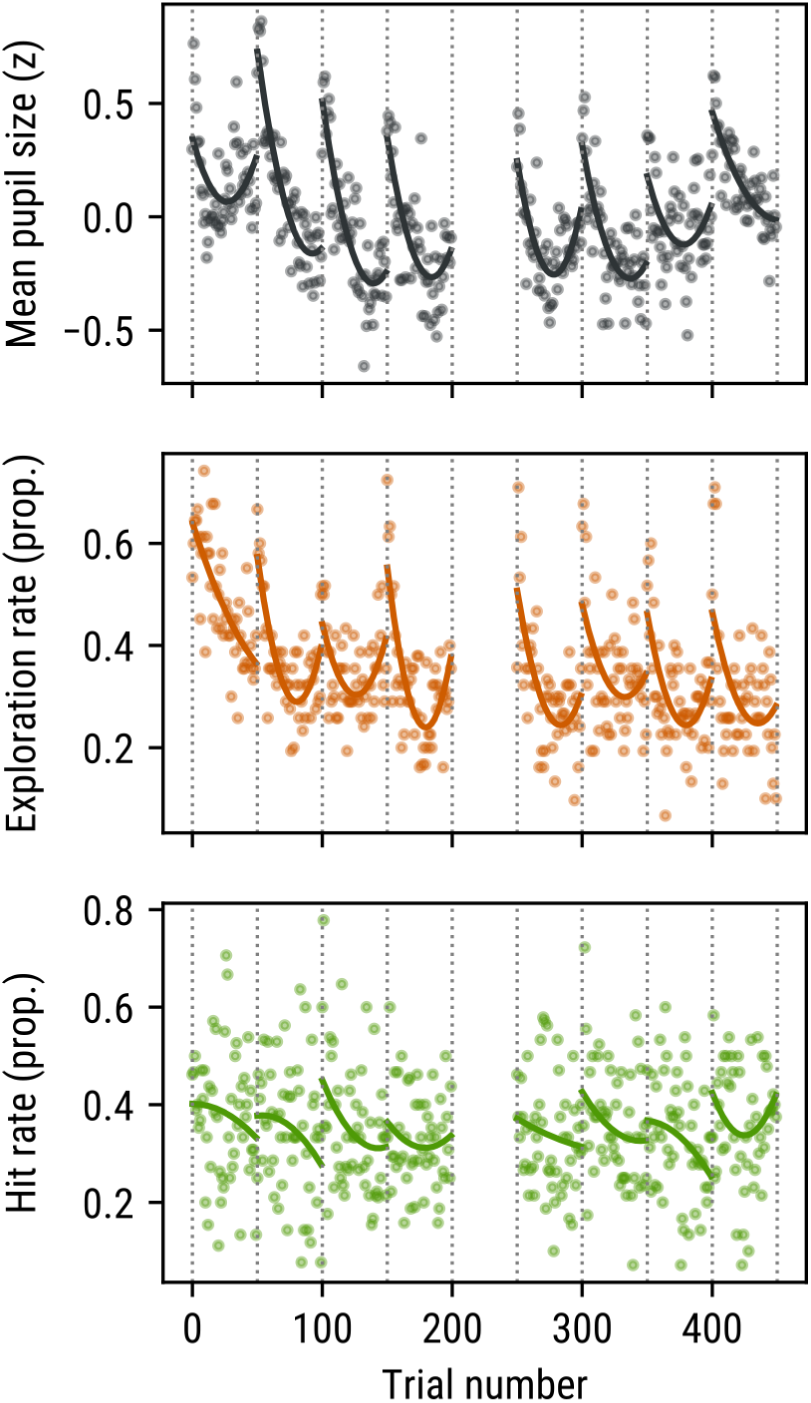
Pupil size, exploration rate, and hit rate (target-present trials only) vary systematically over the course of the experiment. Symbols indicate average values. Lines indicate second-order polynomial fits for each block separately. Trials 200 to 249 correspond to a staircase block, which was excluded from the analysis.

The fact that pupil size, exploration rate, and hit rate are all affected by qualitatively similar time-on-task effects introduces correlations that account for at least some of the effects that we will describe below. Therefore, we calculated residual measures of pupil size and exploration status from which time-on-task effects are removed. Specifically, for each block separately, we fit a second-order polynomial with trial number as predictor and pupil size or exploration status as dependent variable. We then used this fit to subtract predicted pupil size or exploration status on each trial from the observed value. Several of the analyses below are also performed on the resulting residuals, which are not affected by time-on-task effects.

### Exploration is associated with larger pupil size

We ran a linear mixed effects model with pupil size as dependent measure, exploration status (exploration vs exploitation) as fixed effect, and by-participant random intercepts and slopes. This revealed a main effect of exploration status, such that pupil size was larger for exploration as compared to exploitation trials (*z* = 2.687, *p* = .007; Fig. 1b). This effect was still present, though reduced, when analyzing residual pupil size and exploration status (*z* = 2.131, *p* = .033; no random slopes due to convergence error).

Our measure of exploration behavior is based on a mathematical model of decision making. However, a simpler measure of exploration behavior is simply the number of times that participants select the same bandit in a row. A repetition value of 0 corresponds to exploration, while higher repetition values correspond to increasing levels of exploitation. We therefore ran a linear mixed effects model with pupil size as dependent measure, bandit repetition (capped at 3 to avoid extreme values) as fixed effect, and by-participant random intercepts and slopes. This revealed a main effect of bandit repetition, such that pupil size decreased with increasing bandit repetition (*z* = −3.179, *p* = .001; Fig. 1b).

In sum, exploration is associated with larger pupil size. This effect is not fully due to time-on-task correlations, and is robust to different measures of exploration behavior.

### Detection sensitivity is associated with larger pupil size

For target-present trials only, we ran a generalized linear mixed effects model (logit link) with hit status (hit vs miss) as dependent measure, pupil size as fixed effect, and by-participant random intercepts (no slopes included due to converge errors). This revealed a main effect of pupil size, such that hit rate increased with pupil size (*z* = 5.563, *p* < .001; Fig. 1c). This effect was still present, though again slightly reduced, when analyzing residual pupil size and exploration status (*z* = 4.796, *p* < .001).

In sum, detection sensitivity, operationalized as hit rate on target-present trials, is associated with larger pupil size. This effect is not fully due to time-on-task correlations.

### Exploration-driven pupil dilation does not enhance visual sensitivity (in this paradigm)

We ran a generalized linear mixed effects (logit link) model with hit status (hit vs miss) as dependent measure, exploration status (exploration vs exploitation) as fixed effect, and by-participant random intercepts (no slopes included due to converge errors). This did not reveal a main effect of exploration status (*z* = −1.632, *p* =.103; Fig. 1c). (When analyzing residual exploration status, the effect was even reliably in the opposite direction from our prediction, such that increased residual exploration status was associated with a lower hit rate [*z* = −2.360, *p* = .018]. However, since analyzing residual exploration status is not meaningful when exploration status by itself has no effect, we refrain from interpreting this result.)

In sum, exploration-driven pupil dilation does not enhance visual sensitivity, at least not in this paradigm.

## Discussion

The current study investigated the possible perceptual benefits of the pupil-size increase that is associated with exploration behavior. Overall, we successfully replicated two key findings from previous studies: larger pupils were found during exploration trials as compared to exploitation trials (Jepma & Nieuwenhuis, 2011); and larger pupils were associated with an increased detection rate (Ruuskanen et al., 2025; Ruuskanen & Mathôt, 2026b). However, we did *not* obtain evidence for our key hypothesis that the pupil-size increase that is associated with exploration behavior translates to increased visual sensitivity.

First, our results provide an important replication of the study by Jepma and Nieuwenhuis (2011), who found that pupils are larger during exploration than during exploitation—a key finding that has inspired many subsequent studies in the field of pupil-linked arousal. In our study, as in theirs, participants performed a four-armed bandit task. Mean payoffs of each of the bandits changed gradually and independently from one another. Expected payoff for each bandit was followed using a mean tracking rule. Whenever participants chose the highest expected value, the choice was labelled as exploitation; if they chose a different machine, this choice was labelled as exploration. The replication of Jepma and Nieuwenhuis (2011) in our study further supports the Adaptive Gain Theory (AGT; Aston-Johnes & Cohen, 2005) where exploration (or a distractible mode of behavior) is associated with an increase in norepinephrine (NE) and, by extension, with an increase in pupil size (Costa & Rudebeck, 2016; Gilzenrat et al., 2010; Joshi et al., 2016; Murphy et al., 2014; Reimer et al., 2016).

Second, we replicated previous results showing that visual sensitivity increases with pupil size (Ruuskanen & Mathôt, 2026b; Ruuskanen et al., 2025; see also Eberhardt et al., 2022; Mathôt & Ivanov, 2019; Wang et al., 2021). Specifically, one some trials a near-threshold stimulus was briefly presented at an unpredictable location after participants selected a bandit. Participants were better able to detect this stimulus when their pupils were large as opposed to small. This likely reflects in part an optical effect; that is, upon pupil dilation, more light enters the eye (Mathôt, 2018), which increases visual sensitivity. In another part, this likely also reflects an effect of arousal; that is, the Yerkes-Dodson (inverted U-shaped) relationship between arousal and accuracy (Yerkes & Dodson, 1908): accuracy would be highest for intermediate arousal and drop down for both too high and too low arousal. In our results, these two factors likely combine to create the following pattern of results: sensitivity is higher for large than for small pupils, but with a slight drop-off for the very largest pupil sizes (reviewed in Ruuskanen & Mathôt, 2026a).

The main goal of our study was to test whether the pupil-size increase that is associated with exploration translates to an increase in visual sensitivity. If so, this would provide a functional explanation for why pupils dilate during exploration, rather than merely noting that it does, as is usually done in the context of the AGT (Mathôt, 2020; Vilotijević & Mathôt, 2024). However, we did not find any reliable difference in visual sensitivity between trials on which participants explored or exploited. Although one can think of multiple reasons to explain the lack of an effect, in our view the simplest explanation is the most likely: the difference in pupil size was not large enough to have an effect. More specifically, the increase in pupil size associated with exploration might not result in a sufficiently large increase in retinal light influx to lead to a behaviorally measurable increase in visual sensitivity. Possibly, a situation that induces more extreme differences between exploration and exploitation, and thus induces correspondingly larger pupil-size differences, would result in measurable differences in visual sensitivity.

We also investigated whether the associations between pupil size on the one hand, and exploration vs exploitation behavior and hit rate on the other hand, reflect systematic changes in behavior over time (i.e. time on task). This is important because pupil size, exploration state, and hit rate all decrease over the course of a block of trials, thus inducing positive correlations between all three variables. Previous studies have, if at all, tried to account for time-on-task effects by including trial number (within block and/ or within the experiment) as a linear covariate (Brink et al., 2016; Ruuskanen & Mathôt, 2026b). However, the effects of time on task are rarely linear. In the present study, we therefore modeled time-on-task effects as second-order polynomials. We found that all results reported above are indeed partly accounted for by time on task. Importantly, however, all results also hold in slightly weaker form when time on task is controlled for. In other words, the fact that large pupils are associated with increased exploration behavior and visual sensitivity is not only due to the fact that, at the start of each block, pupils tend to be large, behavior tends to be exploratory, and visual sensitivity tends to be high. The association appears to also be driven by spontaneous fluctuations in pupil size and behavior that are independent from time on task (Ruuskanen & Mathôt, 2026a).

To conclude, we replicated the finding by Jepma and Nieuwenhuis (2011) that pupil size is larger during exploration than exploitation. Additionally, we replicated the finding by Ruuskanen, Boehler, and Mathôt (2025; see also Ruuskanen & Mathôt, 2026b) that larger pupil sizes are associated with increased visual sensitivity. However, we did not find that pupil dilation during exploration resulted in increased visual sensitivity, most likely because the difference in pupil size between exploration and exploitation, while reliable, was only modest.

## AI acknowledgement

No AI was used in any stage of writing this paper. AI was used to implement some of the analyses, which were subsequently checked by the authors.

